# Cerebral perfusion metrics calculated directly, model-free, from a hypoxia-induced step change in deoxyhemoglobin

**DOI:** 10.1101/2024.03.08.584122

**Authors:** James Duffin, Ece Su Sayin, Olivia Sobczyk, Julien Poublanc, David J. Mikulis, Joseph A. Fisher

**Author notes:** Corresponding Author: James Duffin, Department of Physiology, University of Toronto, Toronto, ON, Canada.

## Abstract

Dynamic susceptibility contrast (DSC) perfusion MRI measures blood flow metrics via application of a BOLD pulse sequence during transit of a gadolinium-based contrast agent (GBCA), We have previously shown that can also be performed using endogenous deoxyhemoglobin (dOHb) as a replacement for GBCA since dOHb also has the necessary paramagnetic properties similar to GBCA ^1^. Conventional analysis with these contrast agents includes deconvolution of an arterial input function (AIF) assuming a kinetic model. However, the immediate decrease in dOHb that occurs during reoxygenation from a transient hypoxia constitutes a step reduction in susceptibility that permits direct, model-free, calculation of relative resting perfusion metrics. The metrics from this step analysis for seven healthy volunteers were compared to those from a conventional analysis of GBCA and dOHb. Voxel-wise maps of mean transit time, and relative cerebral blood flow and cerebral blood volume, had a high spatial congruence for all three analyses and were similar in appearance to published maps. The time course of the T2*-weighted signal step response identifies both the arrival time, the distribution of the step increase in arterial oxygen saturation, SaO_2_, and the voxel filling phase. The analysis of a step change in dOHb during rapid reoxygenation is an alternative way of calculating perfusion metrics directly from measurements of the resulting T2*-weighted signal, avoiding the potential errors from the use of a kinetic model and requirement for selection of an AIF.

## INTRODUCTION

Dynamic susceptibility contrast (DSC) magnetic resonance perfusion imaging interrogates the voxel-wise passage of a bolus of a susceptibility contrast agent such as a Gadolinium-based contrast agent (GBCA) to measure resting cerebral perfusion metrics including relative cerebral blood flow (rCBF), relative cerebral blood volume (rCBV), and mean capillary transit time (MTT). Analysis of GBCA uses a vascular model-based approach requiring the deconvolution of a tissue concentration time series with a measured arterial input function (GBCA AIF analysis) ^2^. The AIF is based on the strength and temporal evolution of a T2*-weighted signal, sampled in an arterial region of interest such as the middle cerebral artery (MCA) ^3–5^ or choroid plexus ^6^ where the voxel has a blood volume close 100%.

Recent reports describe the use of transient hypoxia-induced changes in deoxyhemoglobin (THx- dOHb) as an endogenously-generated T2*-weighted contrast agent ^1,7,8^. Changes in deoxyhemoglobin concentration ([dOHb]) are produced by targeting lung alveolar oxygen partial pressure (PO_2_) via changes in inspired PO_2_ ^1,7–13^. Resting perfusion metrics using THx-dOHb as a contrast agent can be derived by a similar analysis to that used for GBCA, i. e. a vascular model-based approach using an AIF. This analysis (THx-dOHb AIF) treats the concentration of contrast agent in the voxel as the deconvolution of the tissue signal with an AIF. THx-dOHb AIF analysis has been shown to generate resting perfusion metrics that are comparable to those obtained using GBCA AIF analysis in healthy participants ^8^, and patients with steno-occlusive disease ^14^.

A unique advantage of THx-dOHb as a contrast agent is the ability to manipulate the degree of susceptibility contrast, which is proportional to blood oxygen saturation (SaO_2_), via changes in the PO_2_ in breathed gas ^15–17^. The remarkable physiological observation enabling the generation of an immediate increase in blood SaO_2_ is that full re-oxygenation can occur in a *fraction of a second*. Figure 1 demonstrates the functional anatomy that leads to an abrupt exchange of gases between the lung alveolar gas and the pulmonary capillary blood.

**Figure 1:**
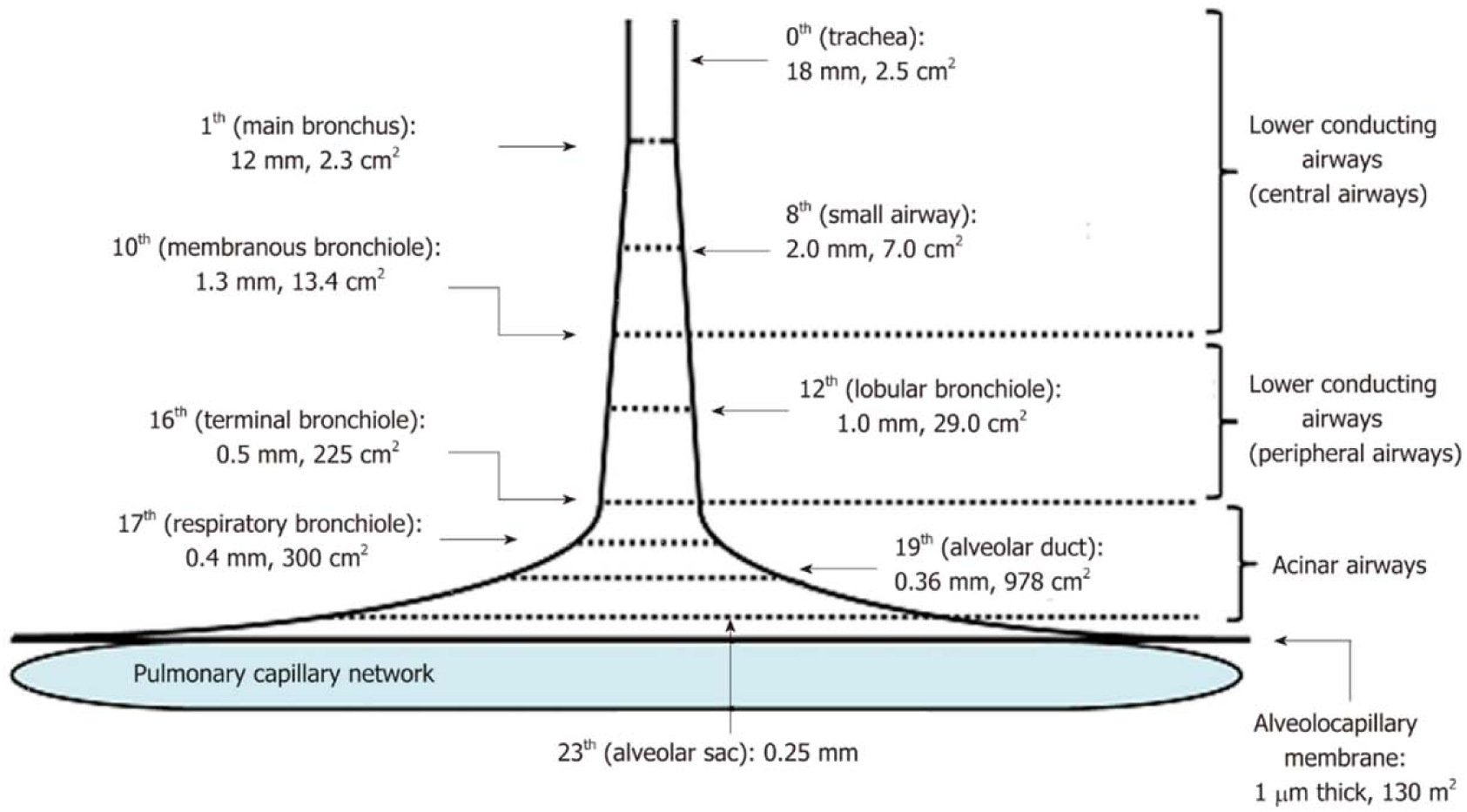
Trumpet or thumbtack model. Schematic presentation of effective diameter and cross-sectional area along lower conducting airways and acinar airways (Figure from Yamaguchi et al 2019 distributed under the Creative Commons Attribution Non-Commercial (CC BY-NC 4.0) license) ^18^.

During an inspiration, inhaled gas passes down successive generations of branches of bronchi and alveolar ducts without undergoing gas exchange. But, in the fraction of a second that the inspired gas passes the 16^th^ branch of airways into the alveolae where gas exchange takes place, the surface area expands from 225 cm^2^ to 130 m^2^ resulting in near instantaneous full gas exchange between air and blood.

Breathing a hypoxic gas can maintain an arterial PO_2_ of 40 mmHg equivalent to an SaO_2_ about 75%, with 25% of hemoglobin in the dOHb state. During a single inspiration of oxygenated air, the pulmonary capillary blood undergoes sudden reoxygenation. The abrupt increased SaO_2_ is conducted into the pulmonary vein, left atrium and ventricle, enters the arterial tree with the same abrupt leading edge of hemoglobin saturation at every branching. This rapid transition from dOHb to oxyhemoglobin describes a step susceptibility input function ^19^. We propose that the resulting increase in the T2*-weighted signal can be used to directly calculate model-free resting perfusion metrics.

The T2*-weighted signal in a voxel is not only dependent on blood volume but also the radius of the vessels in the voxel ^13^. The simplifying assumption for the step analysis of changes in [dOHb] (THx-dOHb Step analysis) is that the T2*-weighted signal (S) in a voxel is inversely proportional to [dOHb] and directly proportional to SaO_2_ ^1,13^. Equation 1 sets out this proportionality.

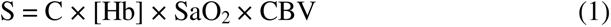

Where:

S = the T2*-weighted signal in a voxel

C = a proportionality constant

CBV = the volume of blood in a voxel

SaO_2_ = the arterial oxygen saturation

[Hb] = the arterial hemoglobin concentration (assumed to be 130 g/L unless measured)

SaO_2_ is related to arterial PO_2_ (PaO_2_) by the in-vivo oxygen dissociation curve ^20^. Equation 2 describes the relation.

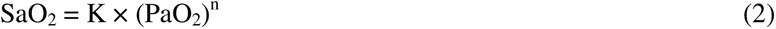

where:

K = 5E-142 × (pH)^157.31^

n = -4.4921 × pH + 36.365

pH is assumed to be 7.4 unless measured.

A step increase in PaO_2_, via reoxygenation, from approximately 40 mmHg to 95 mmHg, produces a step increase in SaO_2_ from 75% to 97%. The step response of the T2*-weighted signal in a voxel increases as blood with saturated hemoglobin displaces blood with deoxygenated hemoglobin. The two models commonly used to depict this displacement process are described in the next section and illustrated in Figure 2.

**Figure 2:**
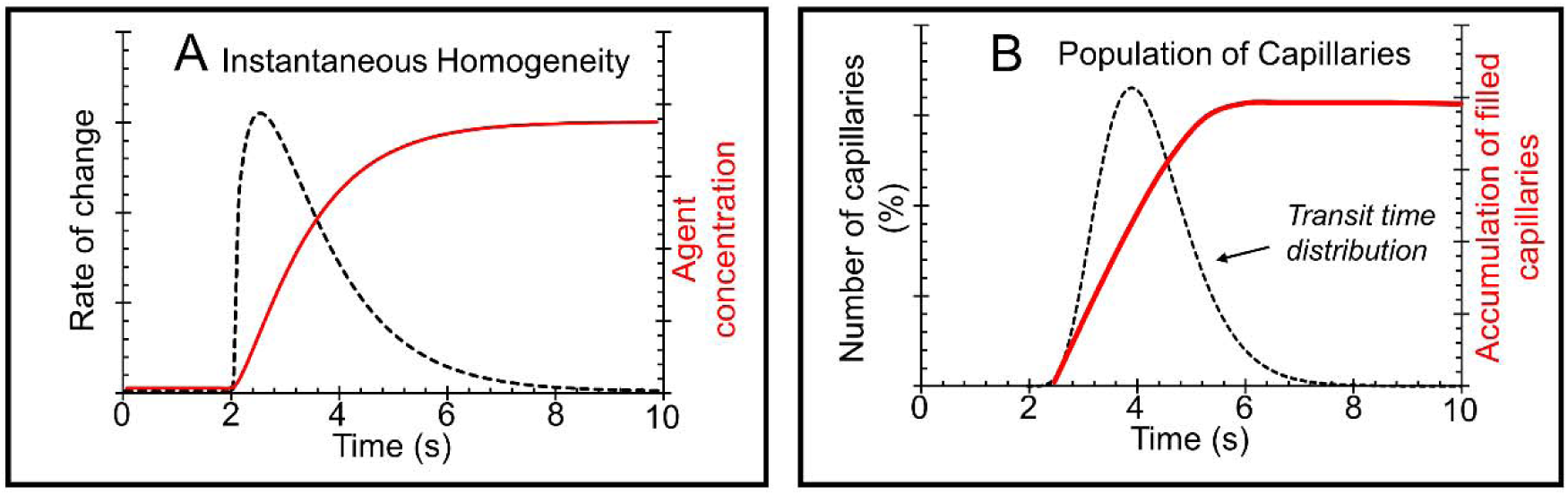
Step response models. A) Instantaneous Homogeneity. Blood with contrast agent enters a well mixed container, so that the initial high rate of increase of voxel contrast agent is a declining exponential shape (dashed line), producing an exponential rise in the voxel contrast agent to an asymptote (red line). B) Population of Capillaries: Blood with contrast agent enters the voxel capillary population, producing an initial linear rise in the contrast agent in the voxel (red line). The distribution of transit times (dashed line) determines the rate of increase of voxel contrast agent, which tapers as capillaries with longer transit times continue to fill.

### Current models for voxel-wise analysis of hemodynamics

If the voxel is viewed as a compartment with instantaneous mixing ^21^ (Figure 2A), the T2*- weighted response depends on the balance of contrast agent inflow and outflow. The initial high rate of T2*-weighted signal increase declines exponentially as the difference in inflow and outflow of contrast declines to zero. The time constant of change is MTT. Thus, model A describes the central volume theorem where MTT = CBV/CBF. This type of kinetic model is assumed for the calculation of perfusion metrics using the GBCA AIF and THx-dOHb AIF analyses.

Alternatively, if the displacement is viewed as the filling of a population of capillaries with varying transit times ^22^, then the rate of increase of the T2*-weighted signal response depends partly on the distribution of capillary transit times (Figure 2B). As blood with the increased SaO_2_ enters the voxel’s population of capillaries, the T2*-weighted signal response increases linearly until capillaries with shorter transit times are filled. Then the T2*-weighted signal rate of increase slows as capillaries with longer transit times complete the filling of the voxel capillary population.

### Direct ‘Step’ analysis

Here we propose to measure perfusion metrics *directly* from measurements of the T2*-weighted signal response to the step increase in SaO_2,_ *without the assumption of either of the kinetic models shown in Figure 2*, the THx-dOHb Step analysis. We assume that equation 1 applies, so that the complete T2*-weighted signal increase resulting from a step increase in SaO_2_ and concomitant decrease in [dOHb], is proportional to the volume of blood in the voxel (rCBV). We demonstrate that this direct examination of the step response generates perfusion metrics and addresses the limitations of the kinetic models shown in Figure 2. Here we compare our proposed THx-dOHb Step analysis of hemodynamic parameters to those generated from GBCA AIF and THx-dOHb AIF analyses using the instantaneous homogeneity kinetic model (Figure 2A), in seven healthy volunteers.

## RESULTS

### Group Comparisons

Figure 3 compares the group averaged metrics from the three analyses in the healthy participants using boxplots, and Figure 4 displays axial slices of the group averaged perfusion metrics and their associated whole brain histograms. While rCBF and rCBV metrics are similar for all three analyses, the MTT for the THx-dOHb is longer that the conventional analysis MTT for reasons to be elaborated upon in the Discussion.

**Figure 3.**
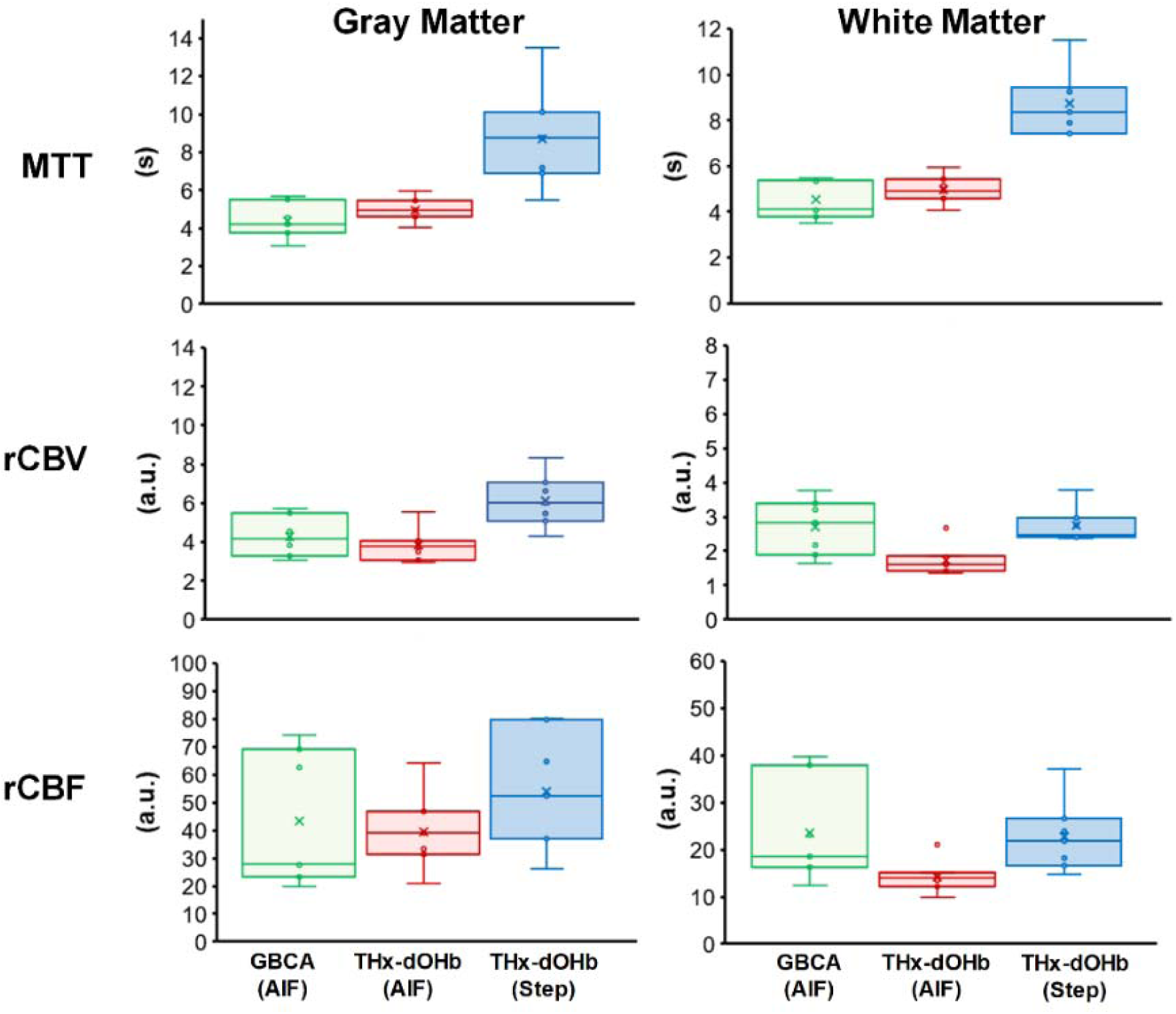
The distribution of the group perfusion metrics in the gray and white matter for the three analyses for the 7 healthy participant group. See table of short forms in the text.

**Figure 4:**
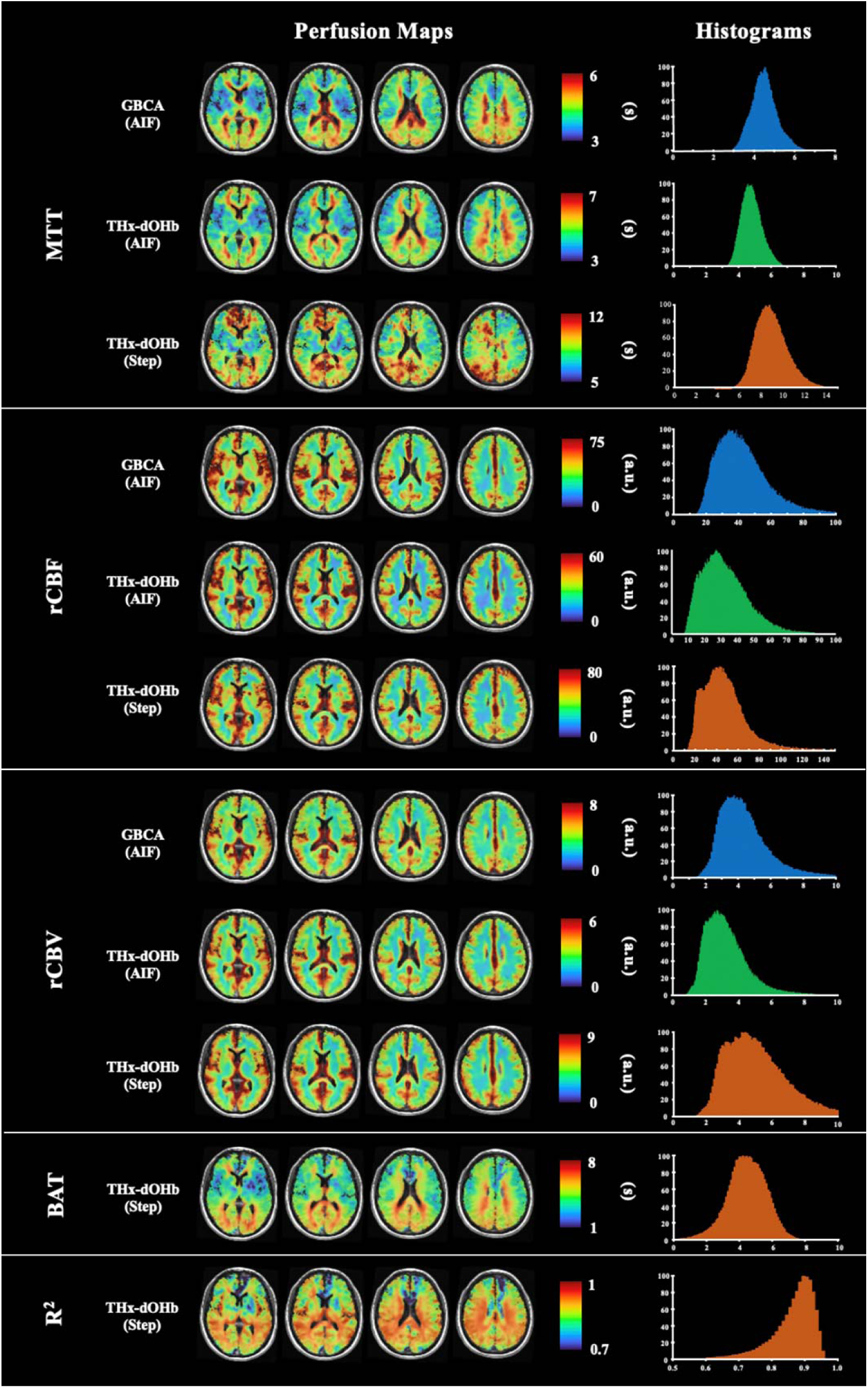
Representative axial slices. The healthy participant group metrics (rCBF, rCBV and MTT) obtained using the three analyses with their associated whole brain histograms. The THx- dOHb Step analysis also provides an R^2^ goodness of fit measure and a blood arrival time relative to the reference cursor time.

### Statistical Results

The rCBV and rCBF measures are relative and expressed in arbitrary units, and so cannot be compared between analyses statistically. However, MTT is in seconds, and a two-way ANOVA was used to examine MTT for differences between analyses and between GM and WM. An All Pairwise Multiple Comparison Procedure (Holm-Sidak method) found no significant interaction between the factors (P = 0.985) and no significant difference between GM and WM (P = 0.667). The Step MTT was significantly greater than the other analyses (including MTT2) (P <0.001).

A one-way ANOVA was used to examine the GM/WM ratios for differences between the 3 analyses. Only rCBF values differed between the analyses, where a Kruskal-Wallis One Way Analysis of Variance on Ranks and an all pairwise multiple comparison procedure (Tukey Test) showed that the GM/WM ratio for the THx-dOHb AIF analysis was higher than that for GBCA AIF analysis (P = 0.02).

## DISCUSSION

This study demonstrates the feasibility a new method of analysis of the T2*-weighted signal increase in response to a step increase in signal resulting from a step reduction in paramagnetic THx-dOHb (THx-dOHb-dOHb Step). The main findings of this study are that the hemodynamic parameters obtained by model-free analysis of THx-dOHb Step were spatially comparable to those obtained with existing kinetic model-based deconvolution analysis which requires an AIF input for both GBCA and THx-dOHb contrast agents.

### Group metrics for rCBF and rCBV

The resting perfusion group metrics for rCBF and rCBV for all three analyses are relative measures. Consequently, comparisons with published absolute data are not possible for numerical values, but only in terms of whether the mapped distributions are realistically like those observed for absolute metrics. The group averaged maps of rCBV and rCBF for all analyses are similar in appearance to each other. Of particular significance, is that they are also strikingly similar in regional distribution to published group average maps of these metrics generated using other analytical methods ^23–27^. However, we note that the rCBV and rCBF maps for the THx-dOHb AIF analysis appear to have lower values than those for both GBCA AIF and THx-dOHb Step analyses. We attribute this finding to a combination of differences in analysis between the THx-dOHb AIF and THx-dOHb Step analyses, discussed below, and the stronger paramagnetic properties of Gadolinium, which may saturate the T2*-weighted signal.

### Group metrics for MTT

MTT for the three analyses provided values in seconds, which can be compared numerically. MTT for the grouped healthy participant maps using GBCA AIF and THx-dOHb AIF analyses depicted regional variations of about 3 – 7 s, whereas MTT values from the THx-dOHb Step analysis are significantly higher. We suggest that the higher values found for the Thx-dOHb Step analysis are the result of measuring MTT directly from the T2*-weighted signal response to the step increase in SaO_2_ as discussed further below.

MTT values from all analyses apart from the step MTT are comparable to published values found for DSC MRI, (mean+/-SD) GM (3.0+/-0.6 s), and WM (4.3+/-0.7 s) by Helenius et al. (2003) ^28^. Using PET measurements of MTT Ibaraki et al. (2007) ^26^ found 3.9 – 4.3 s in the cortex, and using DSC MRI, 2.6 to 3.3 s in GM and 3.5 s in WM. A whole brain average of 5.4 ± 0.92 s was found by Wirestam et al. (2010) ^29^. MTT values from previous experiments using THx-dOHb found values in the 0 - 8 s range ^7^, and 3.9 ± 0.6 s in GM and 5.5 ± 0.6 s in WM ^1^, and 3.5 – 7 s ^8^.

### The step function

The use of rapid conversion of dOHb to oxyhemoglobin to generate a step function has been previously reported in mechanically ventilated rats by Zhao et al. (2021) ^19^ for studying renal hemodynamics. Unfortunately, due to a large apparatus dead space, their reoxygenation took 30 s compared to the 2-3 s achieved here (Figure 7C and Figure 2 in ^1^). Furthermore, Zhao et al. employed oxygen changes from 10% O_2_ (PO_2_ about 75 mmHg) to 100% (PO_2_ about 714 mmHg) for reoxygenation. This high PO_2_ is unnecessary for re-saturation of hemoglobin as hemoglobin is almost fully saturated at a PO_2_ of about 90 mmHg ^20^. Finally, this high PO_2_ in the FRC of the rats after re-oxygenation resists further deoxygenation by dilution which is required for a repeat test. Here, the ability to precisely control inspired PO_2_ has provided the ability to rapidly repeat transient hypoxia and generate a step increase in SaO_2_ during resaturation.

### Models

The GBCA AIF and THx-dOHb AIF analyses assumed a kinetic model with first order dynamics. For a bolus of contrast agent the AIF analysis, via the two models depicted in Figure 2 cannot be distinguished. The instantaneous homogeneity model illustrated for a step change shown in Figure 2A was assumed for the AIF analyses. In that case the residue function is an exponential curve with a time constant MTT as stated in equation 3. The analysis of the T2*- weighted signal change in response to a step increase in SaO_2_ provides evidence to test this assumption. If the SaO_2_ increase is indeed a step change, then the T2*-weighted signal step response should appear as predicted by one of the models shown in Figure 2. However, the T2*- weighted signal step response observed *did not follow either of these patterns*.

This observation suggests that a modified version of the model proposed by Jesperson et al. (2012) ^22^ is suitable as a description of the observed T2*-weighted signal step response, with contrast agent accumulation dynamics in a voxel viewed as the result of a distribution of contrast agent arrival times in a population of capillaries rather than the synchronous arrival assumed in Figure 2B and Figure 1 in Jespersen et al.^22^. With a step increase in SaO_2_, the time course of the T2*-weighted signal step response reflects both the arrival time distribution of the step increase in SaO_2_, and the voxel filling phase. The model is described in Figure 5 and the hemodynamic calculations in Figure 8. Our model assumes that the T2*-weighted signal step response begins as blood with increased SaO_2_ enters the voxel. Although the model assumes a step increase in hemoglobin oxygenation, the arrival time of the saturation front will vary as there is a range of blood flows and path distances to the voxel. Once all vessels entering the voxel have oxygenated blood, the signal will rise linearly. As the *rate of increase* of oxygenated blood is reduced, the rate of signal change will slow and approach zero.

**Figure 5:**
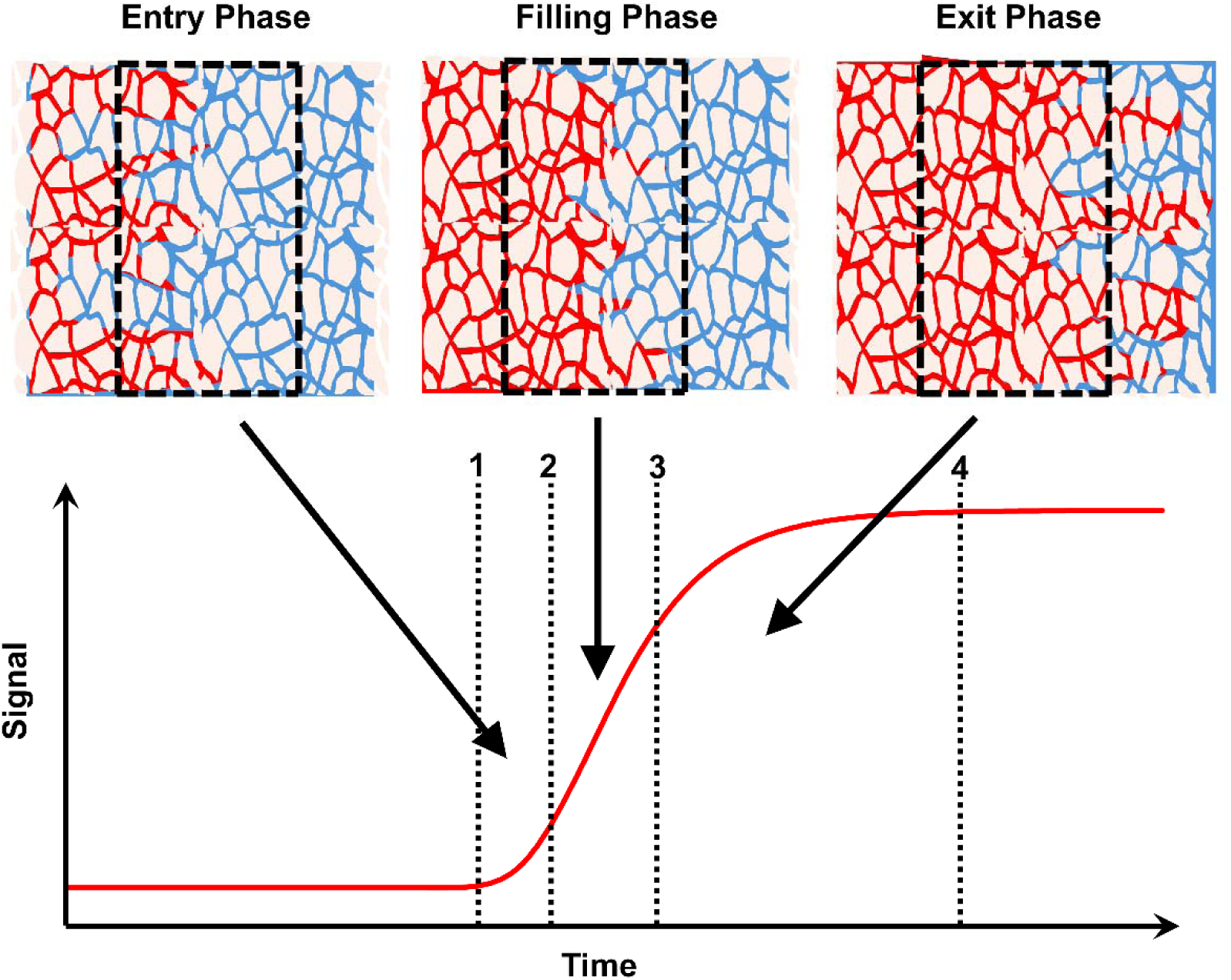
An illustration of the proposed model. The graph shows the wavefront of the step increase in SaO_2_ as it passes through a voxel to fill all vessels with the increased SaO_2_. At line 1 in the graph, the entry phase illustrates the arrival of oxygenated blood (red) displacing the hypoxic blood (blue). Entry is complete at line 2 in the graph and the filling phase illustrates the constant rate of filling with oxygenated blood in all vessels, resulting in a voxel flow of CBF and a linear increase in signal. At line 3 in the graph, the linear stage of filling ends and the exit phase illustrates the slowing of the net rate of increase in signal following a negative exponential pattern, reaching an asymptote of zero change at line 4.

The THx-dOHb Step metrics are calculated directly from the T2*-weighted signal measurements without the assumption of a kinetic model and choice of an AIF. The MTT values are larger than those measured using an AIF and deconvolution because the initial contrast entry time is included in the measurement. In contrast, for AIF and deconvolution analysis the initial entry time is assumed to be the same for all vessels passing through the voxel as illustrated in Figure 2. Indeed, when MTT2 is calculated from the THx-dOHb Step response without the inclusion of the initial entry time constant (c-D in Figure 8), it does not differ from MTT calculated using an AIF, as illustrated in Figure 6.

**Figure 6:**
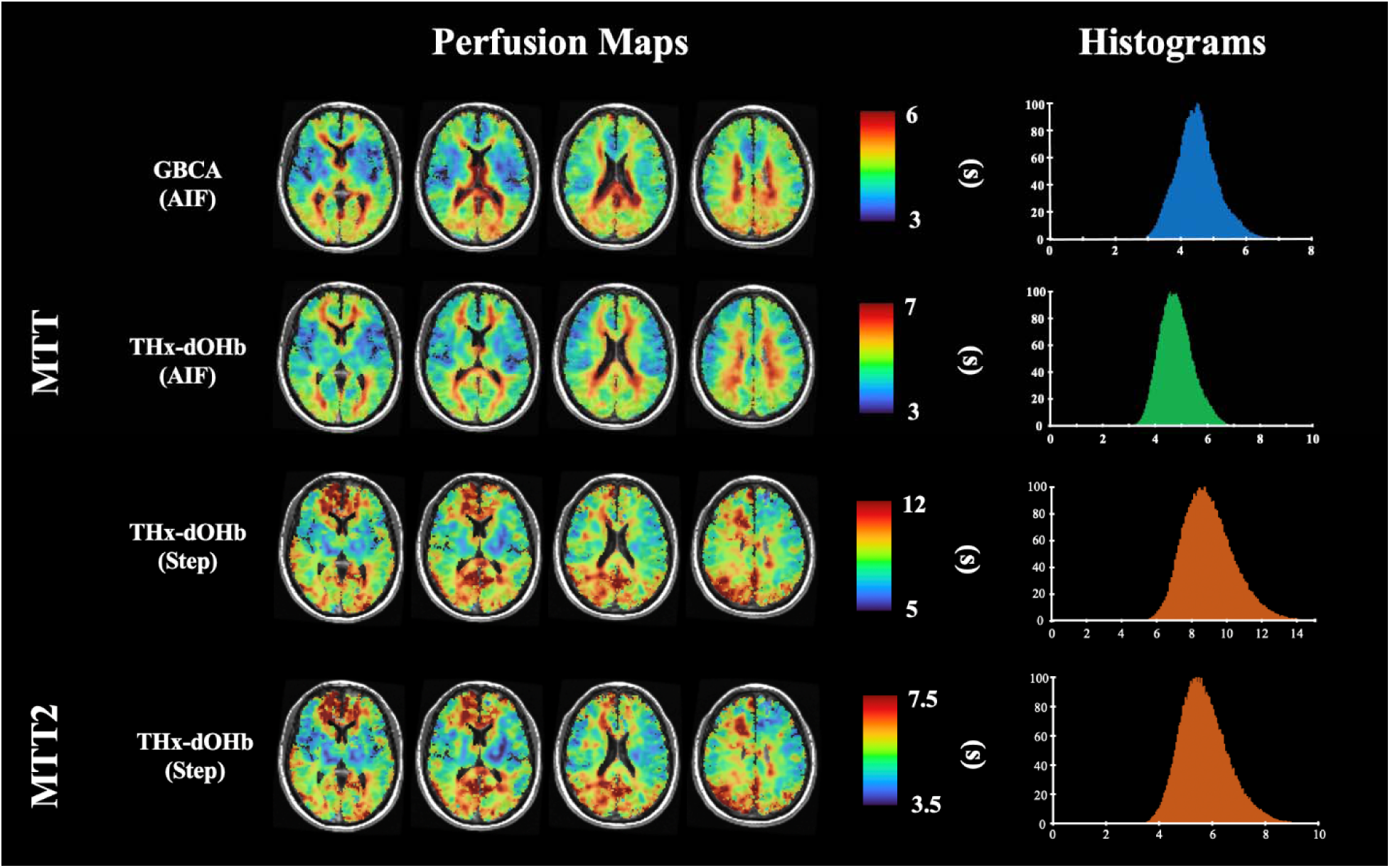
A comparison of MTT maps using all analyses. The healthy participant group MTT metric obtained using the three analyses with their associated whole brain histograms. MTT2 in the fourth row is calculated from the THx-dOHb Step response without the inclusion of the initial entry time constant. The colour scales ranges were adjusted to obtain the highest contrast.

## Study strengths and limitations

The healthy participant group was small and varied in age and sex, so that the results cannot be controlled by either demographic. Data do not reflect the normal range in any specific population. A strength of the study is that we deliberately sampled a wide variety of individuals and focussed on comparison between analyses, whereby everyone was their own control.

We assumed that the transient changes in PaO_2_ do not affect CBF. However, the 60 s exposure to hypoxia may produce an increase in CBF, and act as a confounder for the use of THx-dOHb as a contrast agent, in particular underestimating rCBV and the derived rCBF for the THx-dOHb AIF analysis. Evidence of such an effect is not conclusive (Figure 7). First, there is considerable variation in the CBF response to hypoxia among individuals. The steady-state increase in CBF as PETO_2_ declines from 95 to and 40 mmHg varies from as low as 20% to as high as 50% of resting CBF in healthy individuals ^30^. Second, the CBF response to hypoxia is reported to be slow, whereas we found that the time constant for middle cerebral artery velocity changes measured by trans-cranial Doppler is about 30 s (unpublished data). Optimally, the approach of established baseline hypoxia should be adjusted to be achieved within 30 s. CBF should remain unchanged during the step reoxygenation phase.

**Figure 7:**
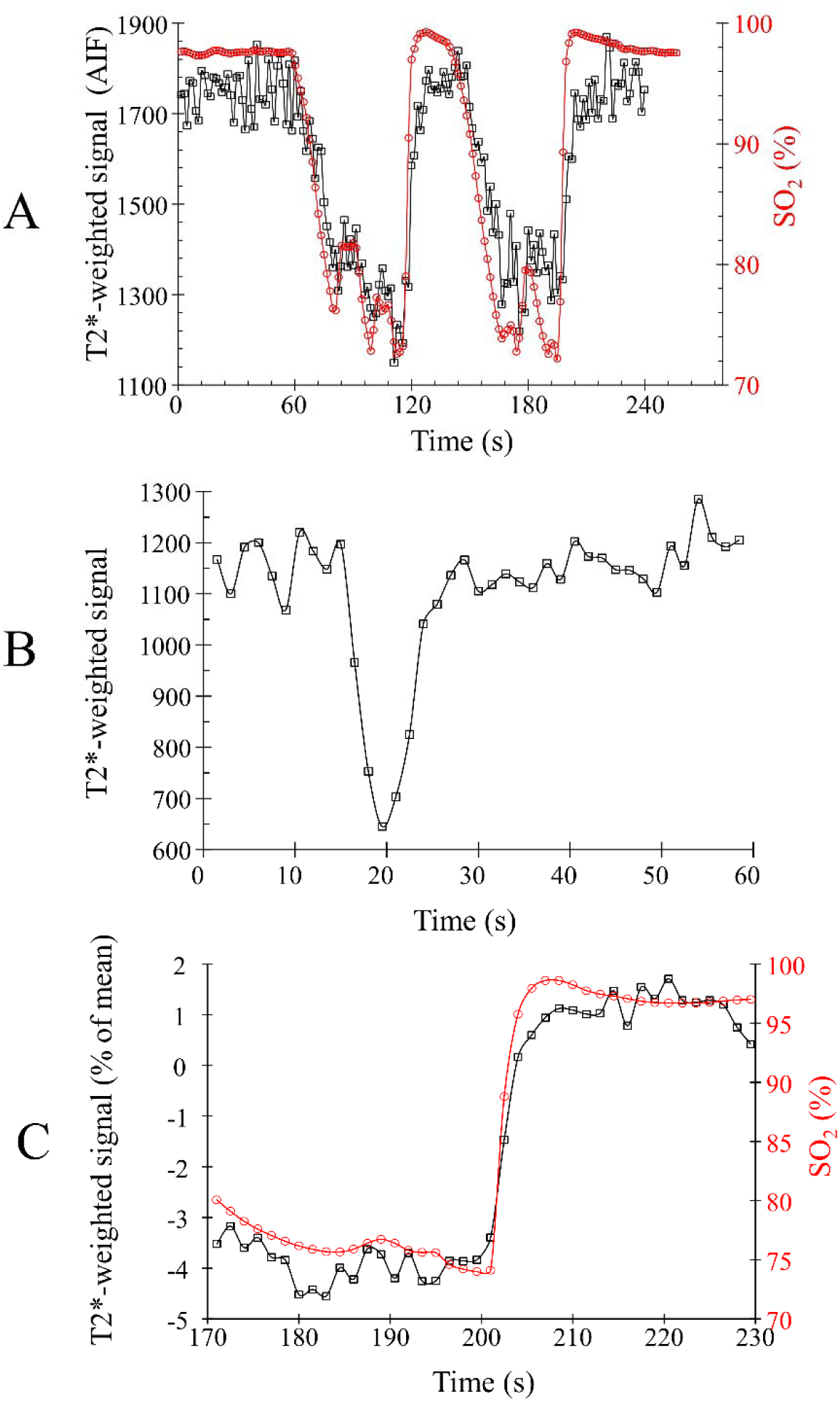
Example data used in each analysis. A) An example of data used for the THx-dOHb AIF analysis showing the hypoxia-induced changes in SO_2_ (%) (red circles) and the AIF T2*- weighted signal (black squares) in a representative participant. B) An example of data used for the GBCA AIF analysis showing the AIF T2*-weighted signal in a representative participant. C) An example of data used for the THx-dOHb Step analysis showing the hypoxia-induced changes in SO_2_ (%) (red circles) and the resulting T2*-weighted signal (black squares) in a representative voxel.

Another assumption underlying these analyses is that the step change in [dOHb] produced in the lungs is preserved as the blood arrives at the cerebral vasculature. We believe this assumption is justified. For a step change in [dOHb] induced in the lungs, any degree of dispersion is less than that observed for an intravenously injected contrast agent. THx-dOHb eliminates an entire pathway over which injected GBCA dispersion can develop, including confluence with contrast-free blood from the brachial vein, subclavian vein, the right atrium, where there is significant contrast dilution and dispersion from non-contrast enhanced blood from the inferior vena cava, the right ventricle, the pulmonary arterial circulation, and the left heart chambers. THx-dOHb dispersion is limited to that produced by passage through the left atrium and ventricle, with the latter dispersion depending on the left ventricular ejection fraction. Normal ejection fractions of about 75%, resulting in roughly one time constant per heartbeat resulting in little dispersion.

The THx-dOHb Step analysis utilises only the step resaturation part of the entire SaO_2_ sequence pictured in Figure 8A and depends on obtaining a rapid step increase in SaO_2_ and a T2*- weighted signal step response free of noise. It is therefore more likely to fail than an analysis of the entire sequence such as transfer function analysis ^12^, which does not require such precise changes in SO_2_, and would be more robust with respect to noise interference. However, the ability to fit a suitable function to the T2*-weighted signal response not only assists with noise rejection, but also enables an increase in temporal resolution and mitigates against such failure.

**Figure 8.**
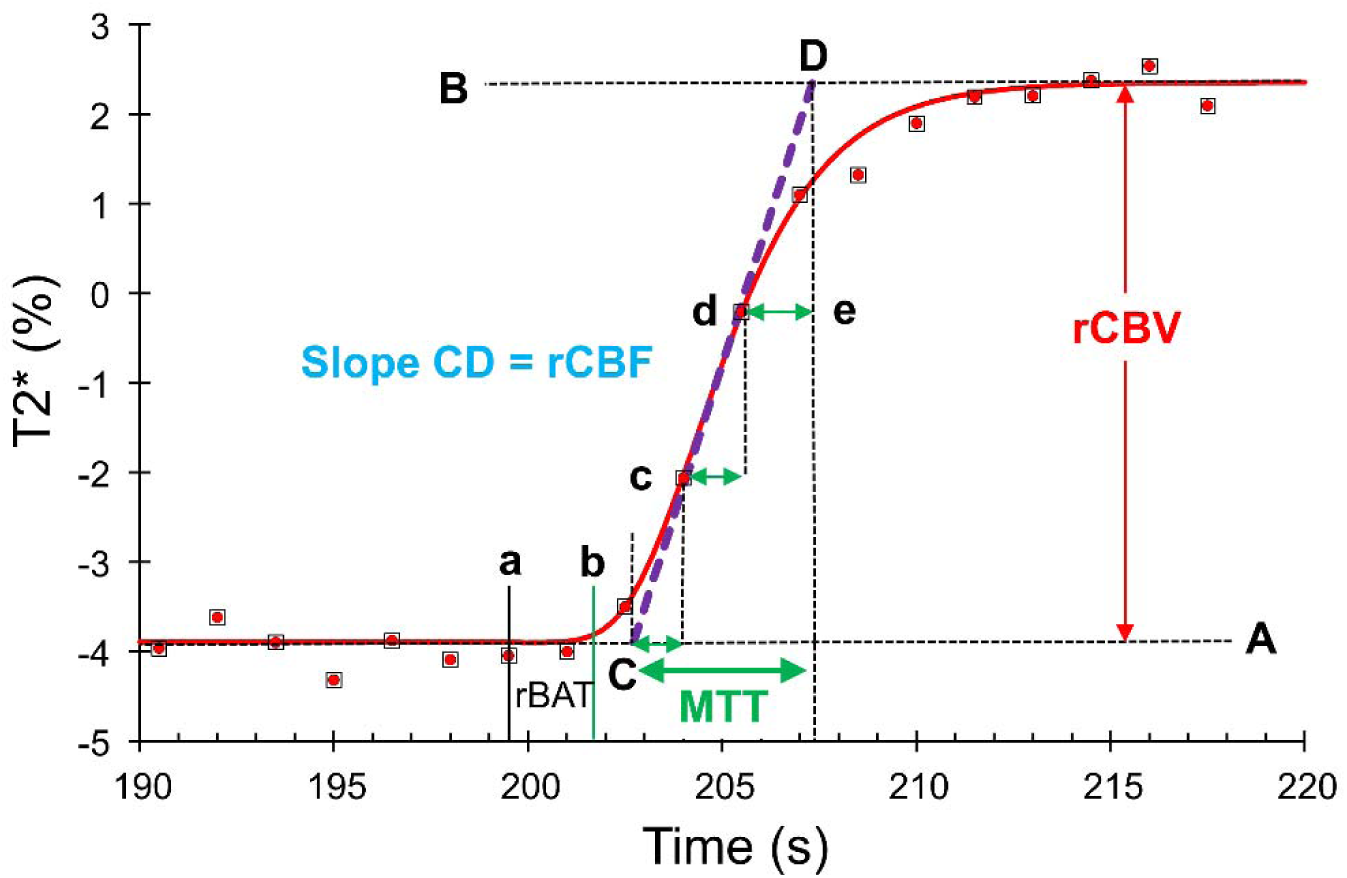
Calculation of hemodynamic variables from voxelwise signal changes following reoxygenation in the lungs. The red filled black squares are the T2*-weighted signal sampled at TR, with Sfit(t) shown as a red line. Then reference time at a defines a-b as the relative blood arrival time (rBAT). C-D is a straight line passing through the linear portion of the signal during reoxygenation (c-d). The slope of this line describes the linear increase in signal reflected by the relative cerebral blood flow (rCBF). The amplitude of the signal change is the relative cerebral blood volume (CBV). The mean transit time (MTT) is the sum of the time constant of voxel filling (C-c), duration of linear filling (c-d) and the time constant of reduction in filling rate (d-D). MTT2 is calculated without the entry time constant as c-D.

We conclude that step reductions in paramagnetic [dOHb] can generate sufficient T2*-weighted signal changes to calculate resting perfusion metrics using a direct model-free analysis. We found perfusion metrics averaged from those of a group of healthy individuals using the step change in SaO_2_ during reoxygenation corresponded strongly, not only with those calculated using a model-based deconvolution of an AIF generated by GBCA and THx-dOHb contrast agents, as well as with other published perfusion metrics. In addition to being non-invasive, the THx-dOHb Step analysis has an advantage over conventional model-based analyses in that it measures perfusion metrics directly from the T2*-weighted signal, model-free, and avoids errors arising from the model assumptions involved in the deconvolution calculations and problematic issues associated with the choice of an AIF.

## METHODS

### Participants and Ethics Approval

This study conformed to the standards set by the latest revision of the Declaration of Helsinki and was approved by the Research Ethics Board of the University Health Network (UHN) and Health Canada. Written informed consent to partake in this study was obtained from all participants. We recruited 7 healthy individuals (5 M) between the ages of 24 and 60 (mean ± SD

= 34.4 ± 16.0 y) by word of mouth. They were non-smokers, not on any medication, and had no known history of neurological or cardiovascular disease.

### Application of Contrast Agents

The standard sequence of changes in SaO_2_ and [dOHb] used for these and other experiments were achieved by controlling the end-tidal partial pressures of oxygen (PETO_2_) and carbon dioxide (PETCO_2_) using the sequential delivery of inspired gases from a computer-controlled gas blender (RespirAct^TM^; Thornhill Medical Inc, Toronto, Canada) running a prospective targeting algorithm ^15^. The principles of operation of the RespirAct^TM^ have been previously described ^31,32^. With this targeting approach, the end tidal values have been shown to be equal, within measurement error, to their respective arterial partial pressures ^16,33^. Reduction in pulmonary PO_2_ is achieved by inhaling successive tidal volumes of prospectively blended hypoxic gas to dilute the oxygen in the functional residual capacity to target a PETO_2_ of 40 mmHg. The re- oxygenation step targets the previous baseline PETO_2_ of 95 mmHg. We avoid overshooting the previous baseline PETO_2_ on re-oxygenation to enable rapid repeats of similar hypoxia-reoxygenation steps.

Participants breathed through a facemask sealed to the face with skin tape (Tegaderm, 3M, Saint Paul, MN, U.S.A.) to exclude all but system-supplied gas. The programmed PETO_2_ stimulus consisted of 60 s at 95 mmHg (normoxia), followed by 60 s at 40 mmHg (hypoxia), 20 s at normoxia, followed by 60 s hypoxia and return to normoxia for 60 s (Figure 7A). PETCO_2_ was held constant at the individual’s resting value. The entire protocol was used for the THx-dOHb AIF analysis, but only the terminal step increase in SaO_2_ was used for the THx-dOHb Step analysis (Figure 7C). SaO_2_ was calculated from the measured PETO_2_ and PETCO_2_, using the oxyhemoglobin dissociation curve relationship described previously in equation 2, assuming a normal pH of 7.4 and a hemoglobin concentration of 130 g/L ^20^. After the completion of the PETO_2_ targeting protocol, the participant returned to free breathing of room air for at least 5 min before an intravenous injection of 5 ml at 5 ml/s of Gadovist (Bayer, Canada), with a baseline delay of 20 s prior to injection, and flushed with 30 ml of saline to be used for the GBCA AIF analysis (Figure 7B).

### MRI Scanning Protocol

The experiments were performed in a 3-Tesla scanner (HDx Signa platform, GE healthcare, Milwaukee, WI, USA) with an 8-channel head coil. The scanning protocol consisted of a high-resolution T1-weighted scan followed by two T2*-weighted acquisition. The high-resolution T1 scan was acquired using a 3D inversion prepared spoiled fast gradient echo sequence with the following parameters: TI = 450 ms, TR 7.88 ms, TE = 3 ms, flip angle = 12^◦^, voxel size = 0.859 × 0.859 × 1 mm, matrix size = 256 × 256, 146 slices, field of view = 220 mm, no interslice gap. Both THx-dOHb and GBCA data acquisitions used a T2*-weighted gradient echoplanar imaging sequence with the following parameters: TR = 1500 ms, TE = 30 ms, flip angle = 73^◦^, 29 slices voxel size = 3 mm isotropic with 64 × 64 matrix.

### Data Analysis

The acquired T2*-weighted signal images were volume registered, slice-time corrected and co-registered to the anatomical images using AFNI software (National Institutes of Health, Bethesda, Maryland) ^34^. The T2*-weighted signal data were converted to a percent of mean. The acquisitions obtained during both THx-dOHb and GBCA were pre-processed in an identical manner to ensure no bias towards any one scan. The ‘spikes’ from the dataset were removed and a spatial blur of 7 mm was applied to each dataset using AFNI software.

First, the images from the THx-dOHb and GBCA acquisitions were analyzed using a conventional kinetic model-based approach ^1^. The visibly sharpest signal change over a middle cerebral artery was selected as the AIF and a deconvolution-based kinetic model of the type shown in Figure 2A was used to calculate voxel-wise maps of MTT and rCBV. A linear relationship between the T2*-weighted signal (S) and [dOHb] is assumed (Equation 1). Standard tracer kinetic modeling was used to calculate MTT and rCBV as stated in equation 3.

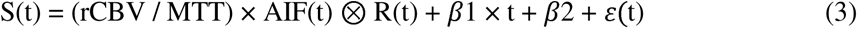

where:

t = time

*β*1 and *β*2 account for linear signal drift and baseline t respectively

ε(t) represents the residuals.

SaO_2_ (t) was used as the arterial input function (AIFt).

R(t) = e^-(1/MTT)^ the residue function (Figure 2A)

The residue function was set equal to 1 at time 0 and 0 at time equal to 5× MTT and bound between 1 and 8 s. Metrics rCBV and MTT were determined using a least square fitting procedure. rCBF was then calculated as the ratio rCBV/MTT using the central volume theorem, (Figure 2A). Values of rCBV and rCBF were respectively multiplied by 7.5 and 30 to obtain easily readable values within the range of absolutes measures.

Second, the voxel-wise analysis of the T2*-weighted acquisition during the THx-dOHb step protocol was implemented using a custom analysis program (LabVIEW, National Instruments, Texas). As Figure 3C illustrates, the step change in arterial PO_2_, via reoxygenation from a hypoxic PETO_2_ of approximately 40 mmHg to 95 mmHg, produces a step increase in SaO_2_, and consequently a step decrease in [dOHb]. The T2*-weighted signal in a voxel increases as blood at the increased SaO_2_ displaces that at the hypoxic SaO_2_. A reference cursor is placed by eye where the whole brain average T2*-weighted signal begins to increase in response to the step increase in SaO_2_ and acts as a time reference for all voxels.

The THx-dOHb Step analysis, illustrated in Figure 8, proceeds as follows: For each voxel, a selected portion of the T2*-weighted signal response before and after the reference cursor (Figure 8, red dots in black squares), is fitted with the Gompertz function specified in equation 4 (Figure 8, red line) using the Levenburg-Marquardt algorithm (LabVIEW module), with R-squared indicating the goodness of fit. Fitting a function to the observed T2*-weighted signal step response served to overcome the inherent noise of the T2*-weighted signal. The Gompertz function, rather than a simple sigmoid function was chosen because it better describes the observed T2*-weighted signal step response.

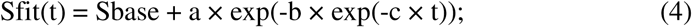

Where

S = T2*-weighted signal

t = time

Sfit(t) = the fitted T2*-weighted signal time course of the step response.

exp = power of e

Fit parameter Sbase sets the initial value of Sfit(t).

Fit parameter a sets the magnitude of the S increase.

Fit parameter b sets the displacement along the time axis.

Fit parameter c sets the rate of change.

Perfusion metrics rCBV, rCBF, MTT and rBAT are all calculated independently as shown in Figure 8 from the Gompertz function fit to the T2*-weighted signal step response, Sfit(t).

Gompertz function fit parameter “a” measures the complete increase in the T2*-weighted step response to calculate rCBV. The start of the T2*-weighted increase (Figure 8, green vertical line) identifies the time where Sfit(t) begins to increase above baseline to exceed 2% of rCBV. Relative blood arrival time, rBAT, is calculated as the difference of start time (b) – reference time (a), with negative values signifying earlier arrival.

The maximum rate of increase of the T2*-weighted signal step response is calculated from the Sfit(t) parameters as “a × c/e” to measure rCBF. Since rCBF is in arbitrary units of T2*- weighted signal % change per second, it is multiplied by 60 to convert to arbitrary units of T2*- weighted signal % change per minute to match absolute units of ml/minute. A line with this slope is drawn thought this point (Figure 8, purple dashed line). It defines three temporal regions, as indicated by the green arrows in Figure 8. First, the exponential increase in signal as the change in SaO_2_ arrives at the voxel until the change has entered all voxels; second, a linear portion of the signal as the vessels fill with the change in SaO_2_ until the change begins to leave the voxel; third, an exponential decay in signal as the SaO_2_ change leaves the voxel. MTT is the sum of the time constants of the first and third temporal regions plus the time taken in the second linear signal increase temporal region. Consequently, MTT satisfies the central volume theorem as the ratio of CBV/CBF.

The perfusion maps obtained from each analysis were transformed into Montreal Neurological Institute (MNI) space and overlayed onto their respective anatomical images. Analytical processing software, SPM8 (Wellcome Department of Imaging Neuroscience, Institute of Neurology, University College, London, UK), was used to segment the anatomical images (T1 weighted) into whole brain cortical supratentorial gray matter (GM) and white matter (WM). Average resting perfusion metrics using all three analyses were calculated for all participants. The MTT, rCBF and rCBV maps for each healthy participant were compiled together to establish average normative ranges for each of the three analyses. This compilation was performed for each metric and analysis by calculating a voxel-by-voxel means and standard deviations from the co-registered maps in standard space ^35,36^.

### Statistical Analysis

Numerical comparisons of the perfusion metrics between the three analyses were not possible for relative values of rCBV and rCBF expressed in arbitrary units. MTT was compared using a two-way ANOVA with factors analysis and region (GM vs. WM). To assess the spatial contrast of the maps, we compared the GM/WM ratios using a one-way ANOVA. Both Normality Tests (Shapiro-Wilk) and Equal Variance Tests (Brown-Forsythe) were part of the ANOVA, and correction for multiple comparisons were applied by an all pairwise multiple comparison procedure. Significant difference for these tests was taken as P<0.05.

## Acknowledgements

The authors thank the MR technologists at Toronto Western Hospital.

## Author Contributions

JAF conceived the study and JD implemented the analyses. DJM selected and reviewed all subjects for suitability. ESS, OS and JP executed the experiments to acquire the data. JD and ESS analysed the data. JD, ESS and JAF wrote the initial draft of the manuscript. All authors participated in the preparation and revision of the final version of the manuscript.

## Data Availability Statement

De-identified, numerical data can be made available to any qualified researcher upon reasonable request to the corresponding author James Duffin.

## Abbreviations

AIF: arterial input function
ANOVA: analysis of variance
DSC: dynamic susceptibility contrast
FRC: functional residual capacity
GBCA: gadolinium-based contrast agents
MCA: middle cerebral artery
MTT: mean transit time
PCO_2_: partial pressure of carbon dioxide
PO_2_: partial pressure of oxygen
rCBF: relative cerebral blood flow
rCBV: relative cerebral blood volume
SaO_2_: arterial hemoglobin oxygen saturation
THx-dOHb: hypoxia-induced deoxyhemoglobin

